# Transcriptomic analysis of dorsal and ventral subiculum after induction of acute seizures by electric stimulation of the perforant pathway in rats

**DOI:** 10.1101/2021.06.14.447935

**Authors:** Beatriz B. Aoyama, Gabriel G. Zanetti, Elayne V. Dias, Maria C. P. Athié, Iscia Lopes-Cendes, André S. Vieira

**Affiliations:** Department of Structural and Functional Biology, Institute of Biology; University of Campinas (UNICAMP), Campinas, São Paulo, Brazil; Brazilian Institute of Neuroscience and Neurotechnology (BRAINN), Campinas, SP, BRAZIL; Department of Translational, School of Medical Sciences. University of Campinas (UNICAMP), Campinas, São Paulo, Brazil

**Keywords:** Hippocampus, Subiculum, Preconditioning

## Abstract

Preconditioning is a mechanism in which injuries induced by non-lethal hypoxia or seizures trigger cellular resistance to subsequent events. Norwood et al., in a 2010 study, showed that an 8-hour-long period of electrical stimulation of the perforant pathway in rats is required for the induction of hippocampal sclerosis. However, in order to avoid generalized seizures, *status epilepticus* (SE), and death, a state of resistance to seizures must be induced in the hippocampus by a preconditioning paradigm consisting of 2 daily 30-minute stimulation periods. Due to the importance of the subiculum in the hippocampal formation, this study aims to investigate differential gene expression patterns in the dorsal and ventral subiculum using RNA-sequencing, after induction of a preconditioning protocol by electrical stimulation of the perforant pathway. The dorsal (dSub) and ventral (vSub) subiculum regions were collected by laser-microdissection 24 hours after preconditioning protocol induction in rats. RNA sequencing was performed in a Hiseq 4000 platform, reads were aligned using the STAR and DESEq2 statistics package was used to estimate gene expression. We identified 1176 differentially expressed genes comparing control to preconditioned subiculum regions, 204 genes were differentially expressed in dSub and 972 in vSub. The gene ontology enrichment analysis showed that the most significant common enrichment pathway considering up-regulated genes in dSub and vSub was *Cholesterol Biosynthesis*. In contrast, the most significant enrichment pathway considering down-regulated genes in vSub was *Axon guidance*. Our results indicate that preconditioning induces synaptic reorganization, increased cholesterol metabolism, and astrogliosis in both dSub and vSub. Both regions also presented a decrease in glutamatergic transmission, an increase in complement system activation, and increased in GABAergic transmission. The down-regulation of proapoptotic and axon guidance genes in the ventral subiculum suggests that preconditioning induces a neuroprotective environment in this region.

## 1. Introduction

Preconditioning is a mechanism in which injuries induced by non-lethal hypoxia or seizures trigger cellular resistance to subsequent events (Amini et al., 2015; Kelly & McIntyre, 1994; Plamondon et al., 1999; Pohle & Rauca, 1994). This injury tolerance seems to be associated with increased expression of genes involved in anti-apoptotic functions, oxidative-stress and heat shock functions (Kirino, 2002; Stenzel-Poore et al., 2003). Norwood et al., in a 2010 study (Norwood et al., 2010), showed that an 8-hour-long period of electrical stimulation of the perforant pathway in rats is required for the induction of hippocampal sclerosis. However, in order to avoid generalized seizures, *status epilepticus* (SE), and death, a state of resistance to seizures must be induced in the hippocampus. The authors used a preconditioning paradigm, consisting of 2 daily 30-minute stimulation periods, to successfully achieve this state of resistance. These authors suggested a decrease in glutamate release by CA3 and Dentate Gyrus (DG) neurons and an increased inhibition of these cells as possible mechanisms to explain how preconditioning by electrical stimulation of the PP induces resistance to seizures in the hippocampus (Norwood et al., 2010). Therefore, this would prevent electrical activity elicited by PP stimulation from spreading throughout the nervous system, triggering SE. However, the authors did not perform experiments investigating the molecular mechanisms and differentially expressed genes induced after the preconditioning in the hippocampus.

The perforant pathway (PP) consists of projections from the entorhinal cortex to the DG (Menno P. Witter, 2007). Layer II neurons of the entorhinal cortex project to the molecular layer of the Dentate Gyrus that in turn project to the stratum lacunosum moleculare of CA3 and CA2. Layer III entorhinal cortex neurons project to CA1 and subiculum neurons (Menno P. Witter & Moser, 2006). Since subicular pyramidal neurons are the main source of projections to deep layers of the entorhinal cortex (Chrobak et al., 2000), the subiculum has a great influence on the electrical output of the hippocampal formation.

The subiculum connects the hippocampus CA1 to the entorhinal cortex (Somjen, 1995; Stafstrom, 2005) and it has an important function in high amplification of neuronal response, short-term memory (Miyashita, 2004) and spatial memory codification (Sharp & Green, 1994a). Furthermore, the subiculum is anatomically divided into dorsal and ventral subiculum, which have specific characteristics associated with its morphology and functions. The dorsal subiculum processes information of space, movement, and declarative memory. Its neurons receive inputs from dorsal CA1 and the entorhinal cortex layer III neurons, projecting outputs to the mammillary nucleus and presubiculum neurons (O’Mara, 2005; Sharp & Green, 1994b). In comparison, the ventral subiculum is responsible for stress modulation through inhibitory projections to the hypothalamic system (Lowry, 2002; O’Mara, 2005; Sharp & Green, 1994b). Its neurons receive inputs from the ventral CA1 and the entorhinal cortex layer III neurons, projecting outputs to amygdaloid complexes and parasubiculum neurons (Lowry, 2002; M. P. Witter & Groenewegen, 1990).

Therefore, due to the importance of the subiculum in the hippocampal formation and its influence on the preconditioning mechanism, this study investigates differential gene expression patterns in the dorsal and ventral subiculum using RNA-sequencing, after induction of a preconditioning protocol by electrical stimulation of the perforant pathway.

## 2. Methodology

### 2.1 Animals

Three-month-old male *Wistar* rats were used for preconditioning protocol (n=16), of which eight animals were electrically stimulated and eight animals received the placement of the electrodes without electrical stimulation, being characterized as a control group. Animals were housed in a 12 hours light/dark cycle on a ventilated rack with free access to food and water throughout experimentation protocol. All the experiments were performed at the University of Campinas- UNICAMP and the experimental protocol was approved by UNICAMP’s research ethics committee (CEUA 3850-1 protocol) according to accepted ethical practices and legislation regarding animal research in Brazil (Brazilian federal law 11.794 from October 8th, 2008).

### 2.2 Surgery and electric stimulation of the perforant pathway

For electrodes implantation, rats were anesthetized using a gas mixture of isoflurane/oxygen mixture (2%/98% respectively) at 2L/min using an acrylic induction box. The anesthesia was maintained during the entire surgical procedure using a mask adapted to an Angle Two (Leica Microsystems, Wetzlar, Alemanha) stereotaxic apparatus. Subsequently, rats were positioned into the stereotaxic apparatus and the skull was exposed by an incision on the scalp where the Bregma was localized and perforations on bone were performed. Next, two bipolar stainless steel, polyamide covered, 0.125 mm stimulation electrodes (P1tec, Roanoke,VA, USA) were placed into the perforant pathway with the following coordinates: +/−4,5 mm lateral, +7,6 mm posterior, and −3 mm ventral).

Electrical activity was recorded by stainless steel, polyamide covered, 0.25 mm monopolar electrodes (P1tec, Roanoke, VA, USA) placed into the dentate gyrus with the following coordinates: +/−2 mm lateral, − 3 mm posterior, −3,5 mm ventral, and −2 mm lateral. In addition, a reference electrode was positioned above the *dura mater* in the coordinates: −1 mm lateral, +3 mm posterior. The final position of recording and stimulation electrodes was based on the occurrence of population spike evoked by granular neurons of DG (Norwood et al., 2010). Furthermore, two perforations in the spaces between reference, stimulation, and recording electrodes were performed to fix stainless steel screws to hold and stabilize all elements into a dental acrylic cement.

Seven days after the electrodes placement surgery, freely moving awake rats were electrically stimulated in the perforant pathway as previously described (Norwood et al., 2010) for 30-min in two consecutive days. The electric stimulation was performed using a Grass Astro-Med S88 stimulus generator (Grass® Technologies, West Warwick, RI, USA), using, paired pulses, 0.1-ms pulse duration, N interpulse interval of 40 ms, and pulse amplitude of 20 V, and the recordings were performed using a miniature preamplifier and digitizer with 32 channels (Intan Technologies, Los Angeles, CA, USA) and the signal was digitized at 10 kHz.

One day after the last preconditioning stimulation session, rats were deeply anesthetized with an isoflurane/oxygen mixture (2%/98%). Subsequently, they were quickly decapitated, and their brains were immediately removed and froze at −60 °C using n-hexane and dry ice.

### 2.3 Laser microdissection

Frozen brains were processed in a cryostat (Leica Biosystems, Wetzlar, Germany) to obtain 60 μm serial sections throughout the entire hippocampus using PEN membrane-covered slides (Life Technologies®, Thermo Fisher Scientific, Waltham, MA, USA). Subsequently, slides were stained with Cresyl Violet, dehydrated with an ethanol series, and stored at −80 °C. A Palm (Zeiss®, Jena, Germany) system was used to delimitate the dorsal subiculum (dSub) and ventral subiculum (vSub), and the tissue was mechanically collected in separate tubes using a surgical microscope (Zeiss®, Jena, Germany) and ophthalmic forceps. The scheme of the rat brain showing the dorsal and ventral hippocampus is shown in figure 1A, and microdissected regions are shown in figure 1B, and figure 1C.

**Figure 1.**
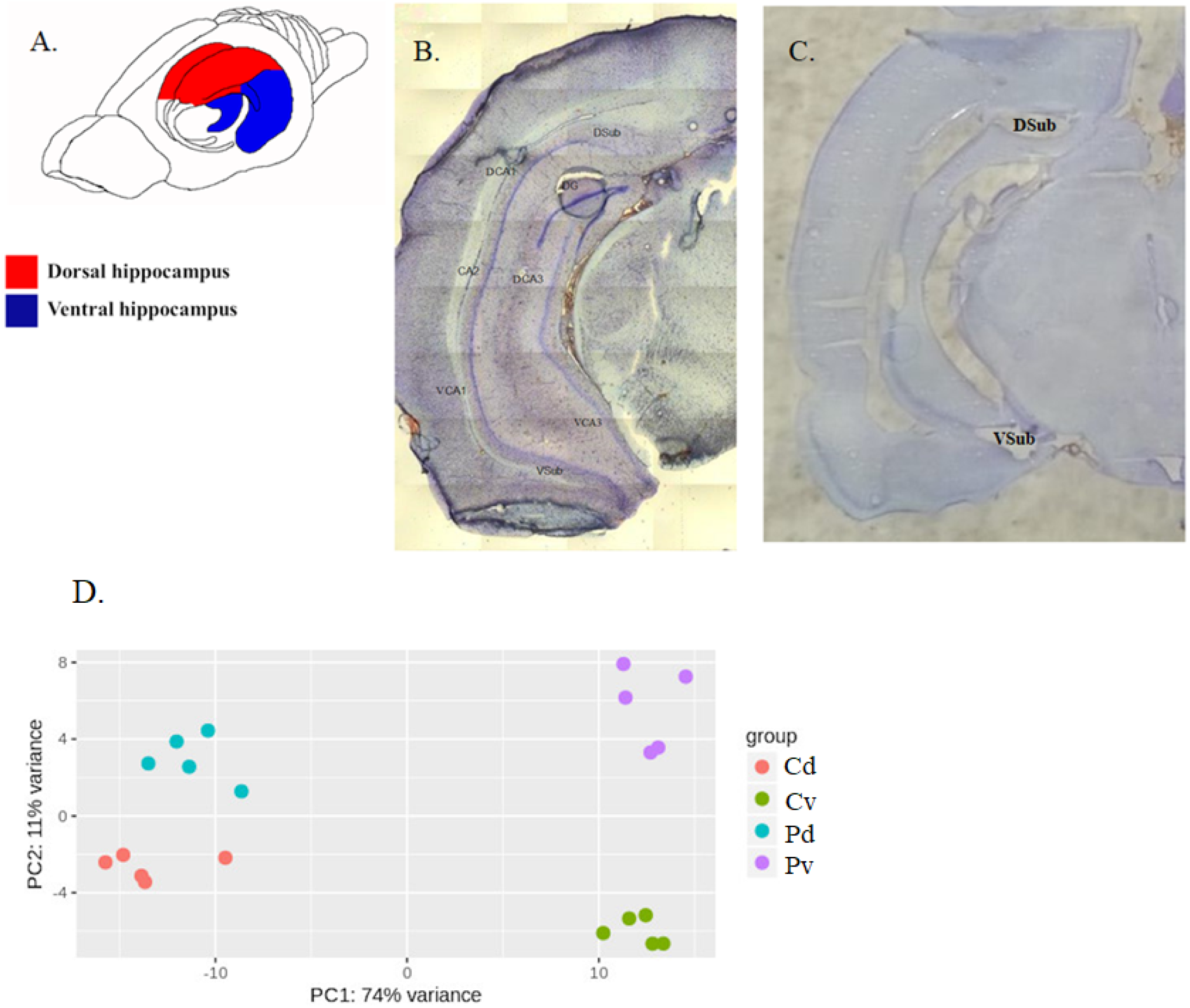
**A.** Adapted scheme of the rat brain showing the dorsal hippocampus in red and the ventral hippocampus in blue. **B and C:** Cresyl Violet stained rat hippocampus section for laser capture microdissection procedures. **B.** The image shows the hippocampal formation structures such as dorsal subiculum (dSub), dorsal CA1 (DCA1), CA2, ventral CA1 (VCA1), ventral subiculum (vSub), dentate gyrus (DG), dorsal CA3 (DCA3), and ventral CA3 (VCA3). The dorsal subiculum (dSub) and ventral subiculum (vSub) were microdissected using LCM, and the tissue was mechanically collected. **C.** Image showing the rat hippocampus after microdissection and removal of dorsal subiculum (dSub) and ventral subiculum (vSub). **D.** PCA graphic for gene expression data. The figure shows the segregation between control and preconditioning groups. In orange is the dorsal control subiculum (Cd), in green is the ventral control subiculum (Cv), in blue is the preconditioning dorsal subiculum (Pd) and in purple is the ventral preconditioning subiculum (Pv).

### 2.4 RNA extraction, cDNA library preparation, and RNA-sequencing

The RNA was isolated from microdissected samples using Trizol’s manufacturer protocol for RNA isolation (Thermo Fisher Scientific, Waltham, MA, USA). Subsequently, the cDNA libraries were produced using the TruSeq Stranded mRNA LT library preparation kit according to manufacturer instructions (Illumina, San Diego, CA USA). Sequencing was performed in an Hiseq 4000 platform (Illumina, San Diego, CA USA) available from Macrogen Inc (Seoul, Korea), producing an average of 17,8 million reads per sample, 95% of bases over Q30.

Reads were aligned to the Rattus norvegicus Ensembl Rnor 5.0 assemble genome using the STAR aligner tool (https://github.com/alexdobin/STAR). The average sequencing alignment rate was 83%. The DESEq2 statistics package (http://www.bioconductor.org/packages/release/bioc/html/DESeq2.html) was used to estimate gene expression and for statistical analysis. A list of differentially expressed genes with a statistical significance of p< 0.05 (after correction for multiple tests) was generated and used for gene ontology analysis with DAVID (https://david.ncifcrf.gov/tools.jsp) and Enrichr (https://amp.pharm.mssm.edu/Enrichr/) software.

### 2.5 Immunolabeling

For immunolabeling, three control and three preconditioning rats were transcardially perfused with 4% formaldehyde 48h after the last stimulation. Brains were processed for histology and immunolabeling for *NeuN* and *GFAP* using fluorescent secondary antibodies. Images were acquired in a confocal microscope (Leica®, Jena, Germany).

## 3. Results

In the present study we found 1176 differentially expressed genes comparing control to preconditioned subiculum regions, in which 204 genes were differentially expressed in dSub and 972 in vSub. Considering the vSub differentially expressed genes in the preconditioned group, 399 genes were down-regulated, and 573 genes were up-regulated (Fig.2A), while in dSub, 71 genes were down-regulated, and 132 genes were up-regulated (Fig.2D).The complete list of differentially expressed genes is presented in the supplementary files. Gene count tables for and raw files (fastq) for all samples are available at GEO (GSE178409).

**Figure 2.**
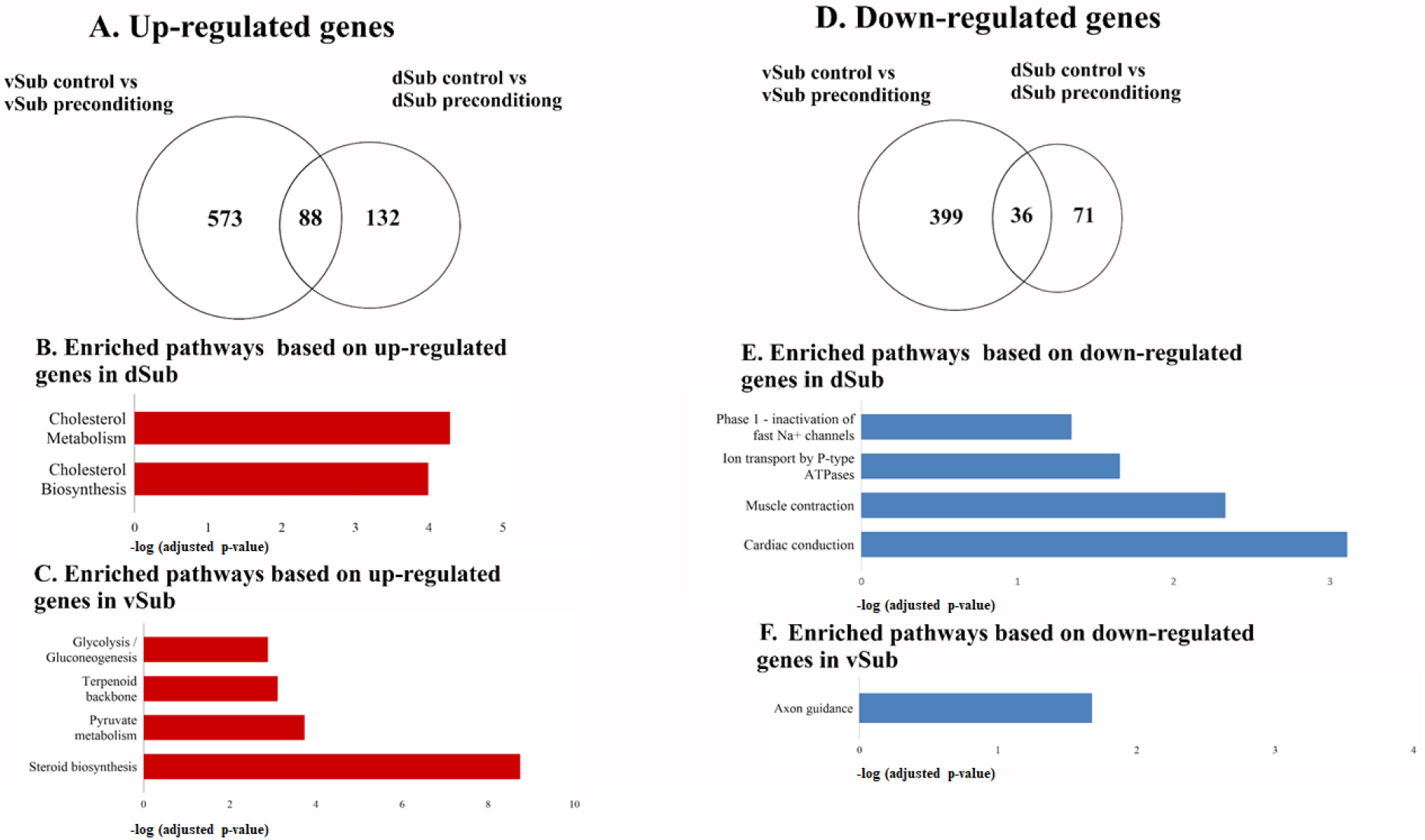
Venn diagram demonstrating the number of differentially expressed genes. The left circle represents differentially expressed genes in the dorsal subiculum (dSub) compared to its control. The right circle represents differentially expressed genes in the ventral subiculum (vSub) compared to its control. The intersection demonstrates differentially expressed genes common to both regions. The graphics show enriched pathways considering differentially expressed genes from dSub and vSub. Moreover, the X-axis represents the ontology pathways, and the Y-axis represents the -log (adjusted p-value). **A.** Venn diagram considering up-regulated genes. **B.** Enriched pathways in the dorsal subiculum (dSub) considering up-regulated genes. **C.** Enriched pathways in the ventral subiculum (vSub) considering up-regulated genes. **D.** Venn diagram considering down-regulated genes. **E.** Enriched pathways in the dorsal subiculum (dSub) considering down-regulated genes. **F.** Enriched pathways in the ventral subiculum (vSub) considering down-regulated genes.

Furthermore, the principal component analysis demonstrated clustering of samples according to the condition, control and preconditioning, in dSub and vSub (Fig.1D).

The most significant common enriched pathways considering dSub up-regulated genes in the preconditioning group were *Cholesterol Biosynthesis* and *Cholesterol metabolism* (Fig. 2B), while in vSub the enriched pathways considering up-regulated genes in the preconditioning group were *Glycolysis/ gluconeogenesis*, *Terpenoid backbone, Pyruvate metabolism,* and *Steroid biosynthesis* (Fig. 2C).

The most significant enriched pathway considering dSub down-regulated genes in the preconditioning group were *Cardiac conduction, Muscle contraction, Ion transport by P-type ATPase, Phase 1-inactivation of fast Na+ channels* (Fig. 2E). In contrast, in vSub the most significantly enriched pathway considering down-regulated genes in the preconditioning group was *Axon guidance* (Fig. 2F). The differentially expressed genes associated with each enriched pathway are listed in table 1. The complete list of enriched pathways is presented in supplementary table 2.

**Table 1.** Differentially expressed genes in dSub and vSub according to the enrichment pathways after induction of preconditioning.

| **Cholesterol Biosynthesis** |  |
| --- | --- |
| Up-regulated exclusively in dSub | FADS2; HMGCS1; ELOVL5; MSMO1; DHCR7; FADS1; CYP51 |
| Up-regulated exclusively in vSub | SQLE; EBP; NSDHL; SC5D; DHCR24; MSMO1; DHCR7; HSD17B7; LSS; CYP51; TM7SF2; FDFT1 |
| Common up-regulated genes in dSub and vSub | MSMO1; DHCR7; CYP51 |
| **Pyruvate metabolism** |  |
| Up-regulated exclusively in vSub | ALDH3A2; LDHA; ACSS2; ALDH2; ME1; ME3; PDHB; ACSS1; ACACA; ACAT2; PCK2 |
| **Terpenoid backbone biosynthesis** |  |
| Up-regulated exclusively in vSub | FDPS; IDI1; MVK; HMGCS1; PMVK; MVD; HMGCR; ACAT2 |
| Glycolysis / Gluconeogenesis |  |
| Up-regulated exclusively in vSub | ACSS2; PGAM1; ENO1; PDHB; ALDH3A2; LDHA; PFKL; ALDH2; ALDH3B1; GALM; ACSS1; PFKP; PCK2 |
| **Gap junction** |  |
| Up-regulated exclusively in vSub | MAP2K1;MAP2K2;HTR2C;ITPR3;TUBB4B;TUBA4A;TUBA1B;TUBB2B;TUBB5;TUBA1A;TUBB2A;TUBB3;MAPK3;TUBA8 |
| **Axon guidance** |  |
| Down-regulated exclusively in vSub | EPHA4; ROCK2; PTCH1; SEMA4C; ILK; SEMA3F; RND1; EFNB3; PARD3; PLXNC1; SLIT2; CAMK2G; MAPK3 |
| **Cardiac conduction** |  |
| Down-regulated exclusively in dSub | KCNJ4; CACNG7; KCNIP4; ATP2A3; FXYD7; ATP1A1; CACNG3 |
| **Muscle contraction** |  |
| Down-regulated exclusively in dSub | KCNJ4; CACNG7; KCNIP4; ATP2A3; FXYD7; ATP1A1; CACNG3 |
| **Ion transport by P-type ATPases** |  |
| Down-regulated exclusively in dSub | ATP2A3; FXYD7; ATP1A1; ATP9B |

Moreover, we found expression of cell population markers *GFAP* (Glial fibrillary acidic protein), *C3* (Complement C3), and *GAD2* (Glutamate Decarboxylase 2) genes up-regulated in dSub and vSub (Fig. 4A, 4B, 4C), the *GLS2* (Glutaminase 2) gene down-regulated in dSub and vSub (Fig. 4D), and the *GAT-1* (Solute Carrier Family 6 Member 1) gene down-regulated in vSub (Fig.4E). Considering the up-regulation of *GFAP* (Fig. 4A), we performed immunolabeling for this gene in coronal tissue sections containing dSub and vSub 48 hours after preconditioning (Fig 3).

**Figure 3.**
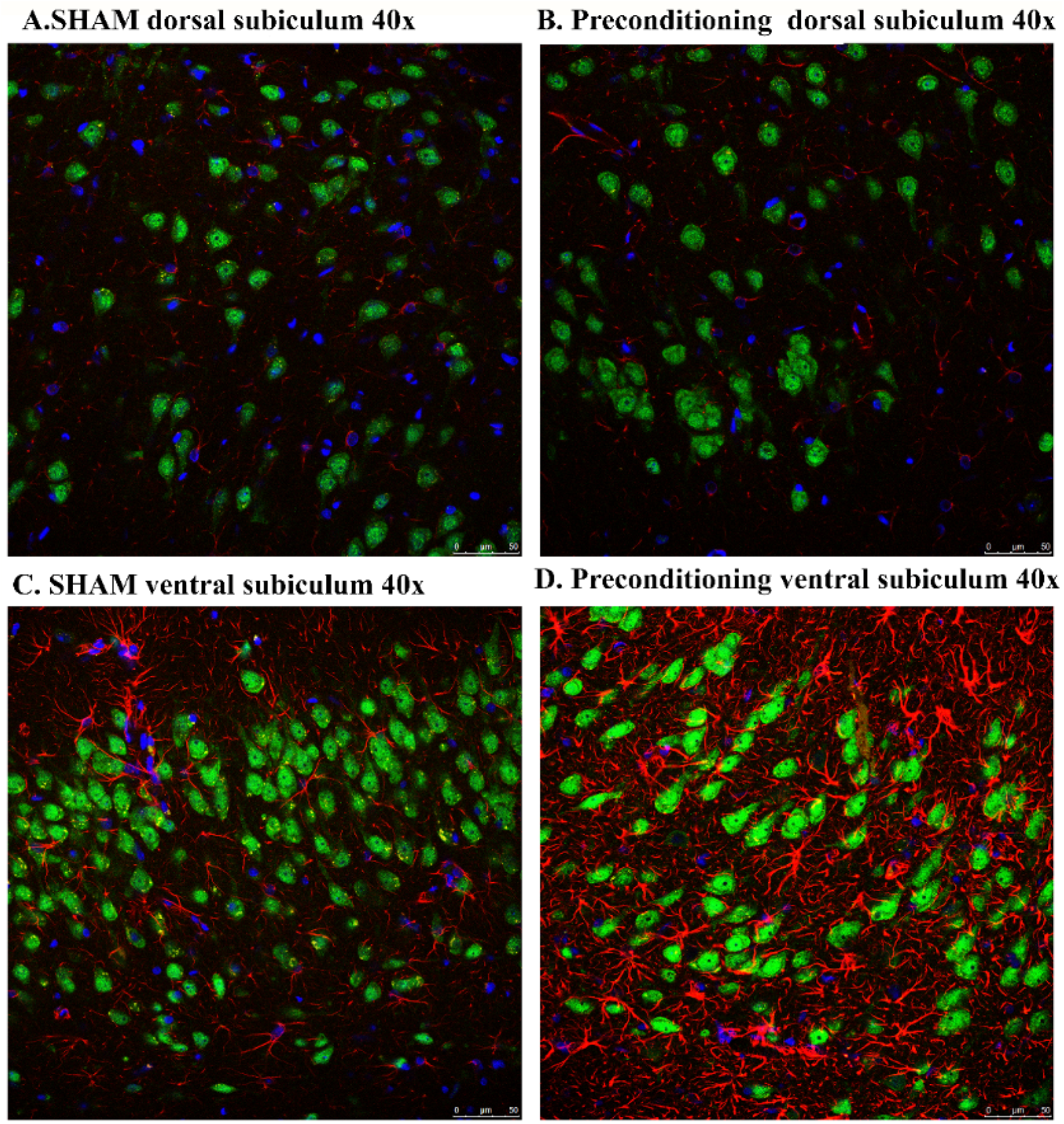
Immunolabeling of the dorsal and ventral subiculum region 48 hours after preconditioning. Scale bar (50μm) at the bottom right of each figure. In green neurons labeled with NeuN, in blue DAPI and in red astrocytes labeled using GFAP. **A.** Control dorsal subiculum. **B.** Preconditioning dorsal subiculum. **C.** Control ventral subiculum. **D.** Preconditioning ventral subiculum.

**Figure 4.**
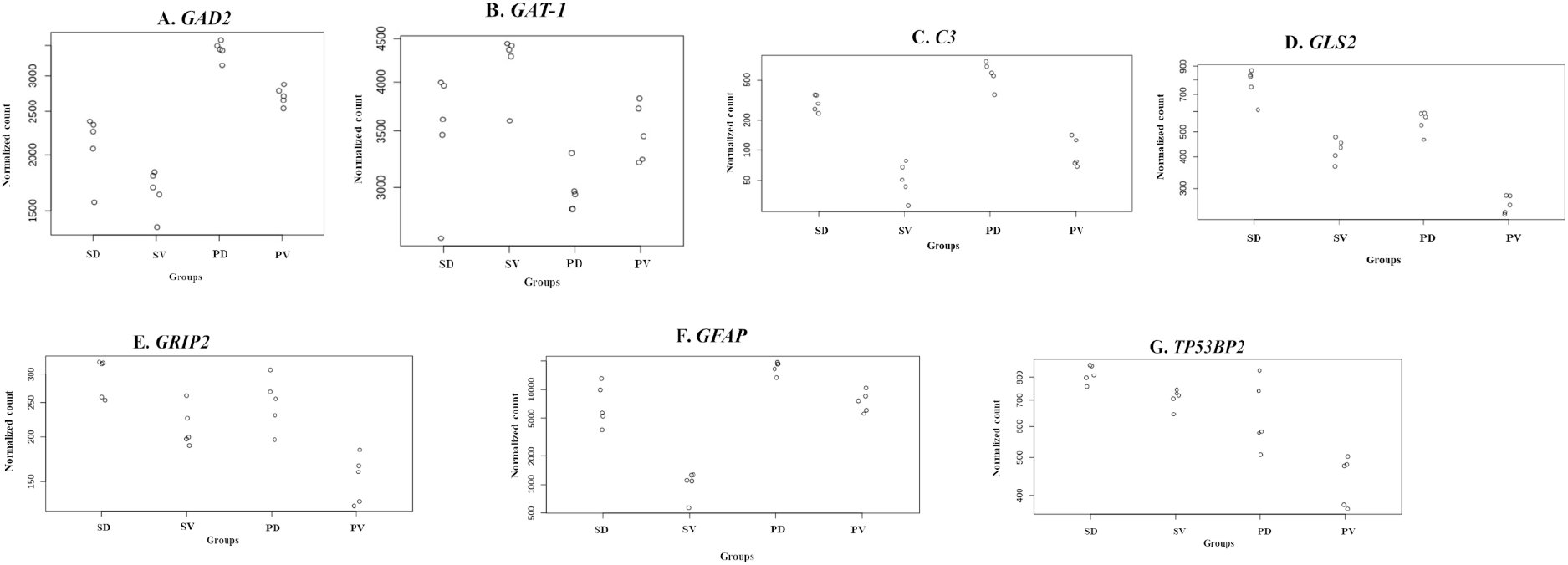
Gene expression estimates based on RNAseq data for GAD2, GAT-1, C3, GLS2, GRIP2, GFAP and TP53BP2 genes. The X axis shows the groups control dorsal subiculum (cd), control ventral subiculum (cv), preconditioning dorsal subiculum (ed), and preconditioning ventral subiculum (ev). The Y axis shows normalized counts for each gene. **A.** Expression of GAD2 gene. **B.** Expression of GAT-1 gene. **C.** Expression of C3. **D.** Expression of GLS2 gene. **E.** Expression of GRIP2 gene. **F.** Expression of GFAP gene. **G.** Expression of TP53BP2 gene.

## 4. Discussion

The present study explores, for the first time, the transcriptome of laser microdissected dorsal and ventral subiculum after a preconditioning paradigm through the induction of acute seizures by electrical stimulation of the Perforant Pathway (PP) in rats. Our results indicate that preconditioning induces synaptic reorganization, increased cholesterol metabolism, and astrogliosis in both the dorsal and ventral subiculum. Both regions also presented a decrease in glutamatergic transmission, an increase in complement system activation, and increased GABAergic transmission. In addition, the down-regulation of proapoptotic and axon guidance genes in the ventral subiculum suggests that preconditioning induces a neuroprotective environment in this region. These altered mechanisms may explain the lack of neuronal loss in the subiculum observed in many MTLE experimental models (Knopp et al., 2005; Stafstrom, 2005).

We observed the up-regulation of GAD2 (Fig. 4A) and down-regulation of GAT-1 (Fig. 4B), suggesting an increase in GABAergic transmission and the down-regulation of GRIP2 and GSL2, suggesting a possible decrease in glutamatergic transmission in both subiculum areas. These findings are in agreement with the hypothesis presented by Norwood et al., 2010 (Norwood et al., 2010) that preconditioning by PP electrical stimulation could result in an increase in inhibition and/or reduction of excitation in the hippocampus. The glutamic acid decarboxylase, GAD2, is present in presynaptic terminals and participates in GABA synthesis. Given its function, the GAD2 plays an important role in central GABA synapses, regulating GABA production (Pan, 2012; Zhao & Gammie, 2014). Thus, the up-regulation of GAD2 gene expression might increase GABA production in presynaptic neurons.

GAT-1 is the most expressed membrane surface transporter of the mammalian brain (Bowery, 2007; Conti et al., 2004). GAT-1 expression is necessary for excitability regulation, synaptic process (Minelli et al., 1995), and reuptake of GABA from the synaptic cleft. The down-regulation of the GAT-1 gene reduces GABA’s reuptake, resulting in enhancement of GABA transmission (Kang et al., 2001). Our data suggest that preconditioning induces increased GABA production by up-regulation of the *GAD2* gene, resulting in increased GABAergic transmission in both subiculum areas. However, in vSub, the GABAergic transmission would be further intensified owing to down-regulation of the *GAT-1* gene, resulting in reduced GABA reuptake.

The present results also revealed up-regulation of the *C3* gene in both subiculum areas after preconditioning (Fig. 4C). C3 is one of the most abundant molecules of the complement system (Stephan et al., 2012). In the CSN, the complement system participates in synaptic pruning (Hua & Smith, 2004; Janeway et al., 2001; Stephan et al., 2012), and in the hippocampus, C3 might be associated with the pruning of glutamatergic synapses (Perez-Alcazar et al., 2014; Salter et al., 2020).

The *GLS2* gene, the glutaminase gene encoding the liver-type isoforms, encodes the GLS2 enzyme, a glutaminase (GA). GA enzymes are considered the main producer of presynaptic glutamate in the brain (Kvamme, 2018; Márquez et al., 2017), participating in glutamate biosynthesis in mitochondria (Suzuki et al., 2010), neurotransmission, and metabolism (Mattson, 2008). The down-regulation of *GLS2* gene suggests reduced glutamate production in dorsal and ventral subiculum neurons. Our data also shows *GRIP2* gene down-regulated (Fig. 4E) in the vSub after the preconditioning. The GRIP2, glutamate receptor interaction binding, is present in excitatory synapses. This receptor binds to glutamate receptors, such as AMPA receptors, inducing their anchoring, trafficking, and recycling (Wyszynski et al., 1999). The study performed by Mao et al., 2010 (Mao et al., 2010; Wyszynski et al., 1999) showed that reduction of GRIP2 could affect the AMPA-R recycling and trafficking, modifying neuronal transmission and synaptic plasticity. The results suggest that down-regulation of GRIP2 might negatively affect the AMPA-R recycling and trafficking, reducing the availability of this receptor on excitatory synapses and reducing excitability in vSub.

The most enriched pathways in both subiculum areas were associated with increased cholesterol biosynthesis (Fig. 2B, 2C). Cholesterol has an important role in the central nervous system (CNS) representing 20-30% of all lipids of the brain (Pfrieger, 2003) and it is locally synthesized by oligodendrocytes, astrocytes, and neurons. In neurons, cholesterol also participates in the production of myelin (Russell et al., 2009), dendritic formation (Fester et al., 2009; Russell et al., 2009); synaptic connections, and axon guidance (de Chaves et al., 1997; Goritz et al., 2005). However, cholesterol production in neurons is inefficient compared to the production in astrocytes (Dietschy, 2009; Nieweg et al., 2009).

Neurons and astrocytes differ in the cholesterol synthesis pathways, neurons mainly use the Kandutsch-Russell pathway, while astrocytes use the Block pathway (Petrov et al., 2016). Our data shows up-regulation of genes participating in cholesterol biosynthesis in neurons and astrocytes. For instance, the gene *NSDHL* participates in the Kandutsch-Russell pathway in neurons and the gene *LSS* participates in the Block pathway in astrocytes. However, most of the differentially expressed genes participate in both pathways, suggesting that increased cholesterol biosynthesis may occur in neurons and astrocytes in both subiculum areas.

Furthermore, RNAseq data showed up-regulation of the GFAP gene (Fig. 4F), and we also found an increase in its immunolabeling in dSub and vSub after preconditioning (Fig. 3). Reactive astrogliosis is characterized by hypertrophy and increase of astrocytes proliferation in injured regions, resulting in protection of new insults (Stevens, 2008), increased neuronal survival (Barreto et al., 2011), and synaptic reorganization (Barker & Ullian, 2010). Therefore, *GFAP* gene up-regulation in dSub and vSub highlights that preconditioning induces astrogliosis in both regions, but mainly in vSub as shown by immunolabeling (Fig. 3C, 3D) and GFAP expression in RNAseq data (Fig. 4A).

Considering down-regulated genes in vSub after preconditioning, our data show an enrichment of the axon guidance pathway (Table 1). Axon guidance is an essential process for the correct function of synapses, influencing positively or negatively the formation of new synapses and remodeling of preexisting synapses (Stoeckli, 2018). Semaphorins, Netrins, Slits, repulsive guidance molecules, and ephrins are proteins expressed throughout axons (Kolodkin & Tessier-Lavigne, 2011; Pasterkamp et al., 2013).

Our data shows the *SEMA3F, SEMA4A, PLXNC1,* and *SLIT2* genes down-regulated (Table 1) in vSub after preconditioning. The *SEMA3F* and *SEMA4C* genes code semaphorin 3F and 4C respectively. The class 3F semaphorins are expressed in the adult nervous system in different cell types, having chemorepulsion or chemoattraction functions (Sahay et al., 2003) depending on the type of receptors they bind. Semaphorin 3F acts as a chemorepulsive signal by binding to the plexin-neuropilin receptor complex, activating signaling cascades depending on focal adhesion kinase (de Castro et al., 1999; Ng et al., 2013); (Kolodkin et al., 1993); (Giger et al., 2000; Huber et al., 2005; Walz et al., 2007). In the nervous system, semaphorin 3F has been shown to inhibit neurite outgrowth and collapse several populations of hippocampus axons (Chédotal et al., 1998); (de Castro et al., 1999). Semaphorins 4A is involved in axon guidance, inducing growth cone collapse in culture and *in vivo* hippocampal neurons (Yukawa et al., 2005) mediated by the Rho/ Rho-Kinase signaling pathway (Jha & Morrison, 2018), promoting neurite outgrowth in cortical neurons (Ishii et al., 2010). Thus, the semaphorins 3F and 4A have a chemorepulsive function in hippocampal neurons.

Our study also found a SLIT2 gene down-regulation in vSub after preconditioning. The SLIT molecule (SLIT1,2 and 3) is a secreted extracellular matrix protein (Holmes et al., 1998) binding with ROBO (ROBO1 and 2), transmembrane glycoproteins of the immunoglobulin superfamily (Skutella & Nitsch, 2001). This interaction allows the collapse of growth cones in axon guidance at Dentate Gyrus and Entorhinal Cortex neurons (Nguyen Ba-Charvet et al., 1999). The down-regulation of SLIT2 may promote axon guidance in vSub after the preconditioning. Thus, the down-regulation of S*ema3F*, *Sema4A,* and *SLIT2* genes may promote axon guidance in vSub after the preconditioning and the increased axon guidance would intensify the GABAergic transmission in the ventral subiculum.

We also found an increase in glycolysis metabolism in the vSub. Glucose is the primary energy source for the nervous system (Jha & Morrison, 2018) being extremely important in the neurotransmission processes. The increased supply of glucose is used for excitatory and inhibitory neurons. Astrocytes take up glucose from the blood capillaries by glucose transporters (GLUTs), and the glucose is stored as glycogen or metabolized into pyruvate in the glycolysis process. The pyruvate is converted to lactate and it is transported to neurons and converted into pyruvate in mitochondria for aerobic energy (Jha & Morrison, 2018). Our results indicate up-regulation of *PFK1, PGAM1, ENO1, LDHA* genes (table 1). These participate in both astrocytes and neuron glycolysis pathways (Yellen, 2018). Thusm suggesting increased energy production might occur in both cells as a possible consequence of the preconditioning stimulation in the ventral subiculum.

Moreover, we observed a down-regulation of *TP53BP2* gene (Fig. 4G) in ventral subiculum. The gene *TP53BP2* encodes the Apoptosis Stimulating Protein of p53–2 (ASPP2) (Iwabuchi et al., 1994; Samuels-Lev et al., 2001; Takahashi et al., 2004) enhancing damage-induced apoptosis (Yang et al., 1999) (Lopez et al., 2000) through the stimulation of p53 transactivation of proapoptotic target genes (Chen et al., 2005). Once the p53 protein is activated, it may induce hypoxic-ischemic and excitotoxic neuronal death (Crumrine et al., 1994). Thus, this gene was associated with proapoptotic functions in neurons, suggesting that its down-regulation might induce a neuroprotective environment in the ventral subiculum, which corroborates with the hypothesis of increased GABAergic transmission in this region. P. The CA3 transcriptome profile of preconditioning animals treated with systemic kainic acid presented down-regulation of genes associated with calcium signalization, ionic channels, excitatory neurotransmitters receptors (Jimenez-Mateos et al., 2008), ubiquitous metabolism, apoptosis and post-translational modifications (Borges et al., 2007). These results suggest the preconditioning neuroprotective role.

One of the goals of this study was to identify the different molecular mechanisms in the dorsal and ventral subiculum after the preconditioning stimulation. Both subiculum areas shared increased astrogliosis (Fig.3), increased GABAergic transmission by *GAD2* gene up-regulation (Fig. 4A), and decrease of glutamatergic transmission by *GLS2* gene down-regulation (Fig. 4D) and *C3* gene up-regulation (Fig. 4C). However, the altered molecular mechanisms were more prominent in the ventral subiculum as demonstrated by the *GFAP* gene up-regulation (Fig. 3 and 4F), the possible increase of GABAergic transmission due to *GAT-1* gene down-regulation (Fig. 4B), and possible decrease of glutamatergic transmission due to *GRIP2* gene down-regulation (Fig. 4E). Only in the ventral subiculum increased expression of genes associated with the glycolysis metabolism pathway (Fig. 2C), down-regulation genes associated with axon guidance (Fig.2F), and *TP53BP2* proapoptotic gene down-regulation were observed (Fig. 4G).

These findings suggest preconditioning by PP electrical stimulations might induce synaptic reorganization, complement system activation, decreased glutamatergic transmission, and increased GABA transmission in subiculum areas. In the ventral subiculum, the down-regulation of proapoptotic genes and axon guidance indicates that preconditioning induces a neuroprotective environment in this region. Although preconditioning induced similar altered mechanisms in both subiculum areas, changes were more noticeable in the ventral subiculum and further studies would be necessary to further explore such region specific effects. Moreover, the results indicate that plastic changes in the subiculum induced by preconditioning by PP electrical stimulation may play a role in regulating hippocampal electrical output during seizures.

## 6. Acknowledgments

We acknowledge the Macrogen Company, Seoul, South Korea, for technical assistance in RNA-sequencing. This work was funded by a grant from Conselho Nacional de Desenvolvimento Científico e Tecnológico, SP, Brazil (CNPq; grant number: 157221/2018-0). Beatriz B. Aoyama is supported by a studentship from CNPq (grant number: 157221/2018-0), Gabriel G. Zanetti is supported by studentship from CAPES, Elayne V. Dias was supported by FAPESP (grant number: 2013/07559-3).

## 8. Author Contributions

**Beatriz B. Aoyama:** Conceptualization, Methodology, Investigation, Validation, Visualization, Formal analysis, Writing—Original Draft. **Gabriel G. Zanetti:** Investigation. **Elayne V. Dias:** Conceptualization, Methodology, Formal analysis, Writing—Original Draft, Supervision. **Maria C. P. Athié:** Conceptualization, Formal analysis, Writing—Original Draft. **Iscia Lopes-Cendes** Conceptualization, Formal analysis, Writing—Original Draft. **André S. Vieira:** Conceptualization, Methodology, Resources, Project administration, Funding acquisition, Supervision, Writing—Original Draft.

## 9. Data Availability Statement

The raw data, the transcriptome data and its analyzes can be disponible upon request to the corresponding author. Gene count tables for and raw files (fastq) for all samples are available at GEO (GSE178409).

